# Brain state dynamics reflect emotion transitions induced by music

**DOI:** 10.1101/2023.03.01.530528

**Authors:** Matthew E. Sachs, Kevin N. Ochsner, Christopher Baldassano

**Affiliations:** Center for Science and Society, Columbia University, New York, NY, USA; Department of Psychology, Columbia University, New York, NY, USA

## Abstract

Our ability to shift from one emotion to the next allows us to adapt our behaviors to a constantly-changing and often uncertain environment. Although previous studies have identified cortical and subcortical regions involved in affective responding, no studies have asked whether and how these regions track and represent transitions *between* different emotional states and modulate their responses based on the recent emotional context. To this end, we commissioned new musical pieces designed to systematically move participants through different emotional states during fMRI. Using a combination of data-driven (Hidden Markov Modeling) and hypothesis-driven methods, we show that spatiotemporal patterns of activation along the temporoparietal axis reflect transitions between music-evoked emotions. Furthermore, self-reported emotions and the subsequent neural response patterns were sensitive to the emotional context in which the music was heard. The findings highlight the role of temporal and parietal brain regions in not only processing low-level auditory signals, but in linking changes in these signals with our on-going, contextually-dependent emotional responses.

## Introduction

At the heart of the word “emotion” lies the Latin word *movere*, meaning to ‘move’. When we suddenly erupt with anger or are swept away by tears, it’s easy to understand the kinetic origins of emotion. Furthermore, our ability to flexibly shift from one emotional state to the next when the situation requires it is important for successful social functioning and wellbeing. Think, for a second, of the social ramifications of not being able to transition from feeling of fear to elation when arriving at a surprise party. And when we do shift to elation, that positive feeling might be experienced quite differently than if we had previously been calmly waiting for friends to arrive rather than startled by friends in the dark. Experimental evidence has confirmed that, in daily life, our emotions reliably transition from one state into another and that these transitions, as well as our predictions for when these transitions might occur, inform our social behavior^1,2^. To date, however, little is known about the neural systems tracking and representing these transitions and the influence of prior emotional contexts. The present paper seeks to address this issue using a combination of fMRI and novel behavioral methods, including the use of music stimuli specifically created for this purpose.

Despite growing behavioral evidence that emotions are dynamic, transitory and situated within a temporal and social context^3^, historically, most neuroimaging studies of emotion have presented participants with static images selected because they elicit specific affective reactions, with analyses seeking to identify the distinct neural bases of different response types rather than understanding transitions between states or the dynamics of emotional responding more generally. While this approach allows for a high degree of experimental control and has conferred new insights into the ways in which the brain processes and represents emotion eliciting stimuli^4,5^, it is less ideal for studying the dynamic changes in states over a longer period of time^6^. Some fMRI studies have used dynamically changing stimuli, such as film clips, to ask how the strength of a given affective state varies in concordance with BOLD signals over time^7–9^. These univariate approaches have shown that various prefrontal and subcortical regions of the brain track the rise and fall of a given emotional state in response to a stimulus (e.g. a film clip) along a prescribed affective continuum (from good to bad or happy to sad). However, they cannot tell us how our brains enable transitions between qualitatively different states nor how one’s recent history of experiencing different states might influence brain functioning. Large-scale behavioral studies have confirmed that the experience of emotion is highly complex and may be best described as a multi-dimensional space^10,11^. A recent study used a data-driven approach to identify distinct neural states that corresponded to emotional responses, but characterized only two broad emotional states and did not assess how state transitions could influence emotional experience^12^. There is therefore a gap between our study of emotions in the lab and the way in which we experience emotions in everyday life, where affective-related information is changing continuously depending on the socioemotional context.

To more reliability and rigorously investigate dynamic transitions between emotional states, unconfounded by other variables like language, we developed a novel musical stimuli set systematically designed to induce a variety of emotional reactions at specific timepoints. For the purposes of studying affective dynamics, music is an ideal stimulus to use in that it is temporal by nature and can reliably express and elicit a range of emotions without language^13^. Despite these methodological advantages, a recent meta-analysis of functional fMRI studies with dynamic and continuously presented stimuli (films, music, speech, video games, etc) found that only 13% of included studies used purely auditory stimuli that did not contain visuals or language^14^. Therefore, it is difficult to determine from these results – or from the vast majority of neuroimaging studies that elicit emotions with visual stimuli – if and how regions in the brain involved in emotional responding are engaged by stimuli whose affect-eliciting properties are abstract and not confounded by changes in language. Knowing this is important not only for assessing the generalizability of our current brain models of emotions, but is relevant for clinical disorders associated with aberrations in non-verbal emotional understanding^15^.

To create this dynamic stimulus set, film score composers were hired to write two pieces of polyphonic, non-lyrical music that convey and transition between 5 pre-defined emotional categories (herein called events). Each piece contained 2-4 events from each emotional category that were musical distinct. Importantly, however, overall instrumentation, tempo, duration and number of transitions were kept constant throughout both pieces. In this way, we could control both the timing of emotional transitions as well as the lower-level acoustic elements that drive these changes. Unlike with previous naturalistic neuroimaging studies that have used pre-existing stimuli, having this degree of control allowed us to more effectively tease apart the different features of the stimulus that are potentially driving the emotional experience and the subsequent brain activation patterns. By commissioning a completely new piece of music, we could additionally minimize the potential confounding effects of familiarity and language and utilize instrumentation and tempos that were most conducive for listening during MRI scanning. While a purely objective “ground truth” may never be realized when it comes to emotions, having access to both the intentions of the composer as well as how listeners perceive, appraise and respond to the music affords us more certainty in knowing which emotions we are studying.

A final advantage of designing a novel stimulus set is that we could systematically manipulate the context in which each event was heard, in order to test the lasting influence of an emotional state on the experience and representation of subsequent states. Previous studies have shown how exposure to an emotional stimulus can bias the way in which new and distinctive information is experienced, learned, and remembered^16^. Using fMRI, it has been shown that the established emotional context in which an event was first encountered influences the neural representation and reactivation of that event^17,18^, particularly in subcortical regions such as the amygdala and hippocampus, as well as cortical regions such as the prefrontal cortex and inferior temporal cortex. When designing the musical stimuli, the composers wrote two versions of each piece, which consisted of the same segments of emotional music arranged in two different ways. We specifically optimized the order so that each event was preceded by a different emotion in each piece. In this way, we can assess how the surrounding context in which an emotional, musical-event is encountered influences how it is experienced and represented in the brain.

Using this novel musical stimulus, fMRI, and a combination of hypothesis-driven and data-driven statistical approaches, we addressed two main research questions. First, we asked which brain regions track emotional state transitions in response to music. For this, we compared voxel pattern stability within vs. across emotional events (hypothesis-driven) and used Hidden Markov models (HMMs) to probabilistically identify brain state transitions (data-driven) without using any timing information about the stimulus itself^19^. In conjunction with dynamic stimuli, HMM-defined shifts in the activity patterns within cortical brain structures (e.g. posterior medial cortex, tempoparietal junction, angular gyrus and inferior frontal cortex, and tempoparietal axis) have been shown to reflect both high level and low-level changes in narrative^16^ and musical structure ^20–22^. Based on these previous findings, we hypothesized that several subcortical (amygdala, putamen, pallidum, and caudate), medial cortical (dorsomedial [DMPFC] and ventromedial prefrontal cortex [VMPFC], and anterior cingulate cortex [ACC]), and temporal lobe (insula, superior temporal sulcus, and PHG) brain regions will show time-varying activation patterns that map onto emotional events in the music.

As part of this first research question, we then further clarify the experiences that might be driving brain state transitions in response to the music. Encoding models were trained to predict fMRI signal based on a weighted combination of musical and acoustical features alone or with the addition of self-reported affective changes in response to the music^23–25^. We predicted that fMRI signal in the VMPFC/OFC, insula and subcortical regions would be better predicted by models that include subjective emotion ratings, in addition to lower level musical/acoustic features.

Second, we asked how the recent emotional context influenced event representations and dynamics. Here, we measured systematic differences in spatial brain patterns associated with an emotional event when it was preceded by different emotional states. Previous studies have used such an approach to show that semantic context and narrative framing can modulate our neural representations of the same stimulus^26,27^. Finally, we tested if the temporal patterns of brain-state transitions are influenced by the nature of the preceding emotion, using state probability metrics derived from HMMs^28^. We hypothesize that regions that are sensitive to emotional transitions in the music will demonstrate faster event transitions when the preceding event was of the same valence. Furthermore, we predict that these regions will show systematic alterations in spatial patterns based on the emotional context in which the same emotional event is heard.

## Results

### Which brain regions track transitions from one emotional state to another?

During fMRI, 38 participants passively listened to two full-length pieces of music (~30 minutes): one version of piece A and one version of piece B, each of which featured 16 emotional events (see Methods and Fig. 1). In between the two music-listening sessions, participants watched a ~12.5-minute audio-visual movie which was used for functional alignment^29^. All results presented below are in shared response space unless otherwise noted.

**Fig 1.**
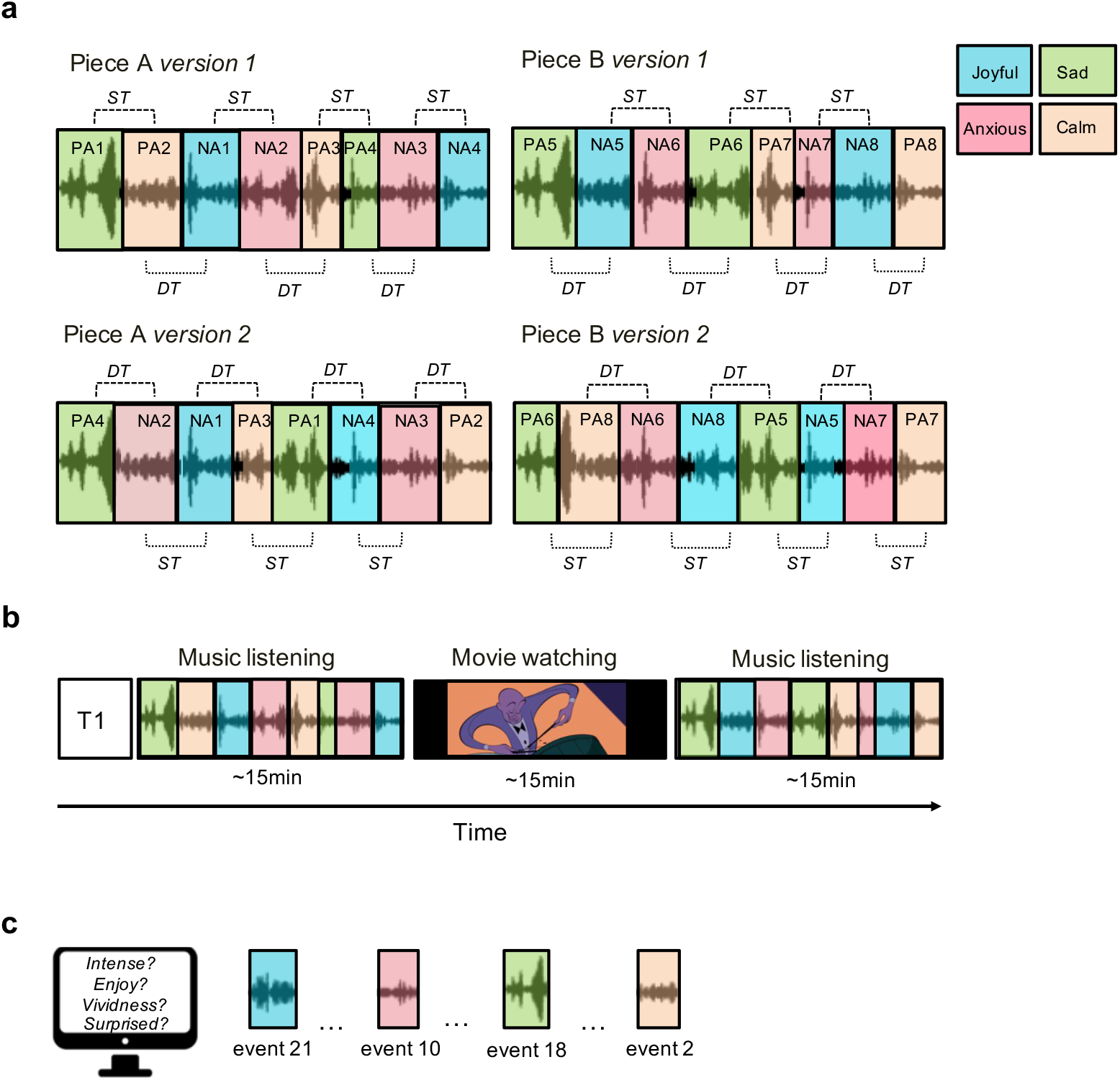
Stimuli and study design. A) Schematic illustration of the two novel musical compositions. Each piece has 16 unique emotional events and two versions. Each version has the same events but in a different order to counterbalance B) Design of the fMRI scanning session in which participants listened to one version of Piece A and B (version randomly selected and counterbalanced across participants). C) Post-scanning recall measures. PA = positive affect, NA = negative affect, PP = positive valence to positive valence emotion transition; ST = same valence transition; Dt = different valence transition.

### Hypothesis-driven approach: within vs. across event temporal correlation results

If a brain region is sensitive to emotional transitions, then we would expect that the brain patterns at timepoints within a particular emotional event should look more similar than brain patterns at timepoints that cross emotional event boundaries. To this end, for each searchlight on the cortical surface of the brain, as well as 11 subcortical ROIs, we computed correlations between all pairs of timepoints (using the functionally-aligned feature space derived through hyperalignment). We then computed the average correlation for timepoint pairs that were within an emotional event (as defined by the composers) and for pairs that spanned two adjacent events (i.e. across event t and an adjacent event t + 1). The difference between the within event correlation and the across adjacent event correlation indicates the extent to which a brain region’s activation pattern shifted at event transitions. Regions that showed significantly higher correlations between time points within an emotion vs. across emotional boundaries included the bilateral auditory cortex, superior temporal and middle temporal gyrus, and temporal pole as well as the left supramarginal gyrus and angular gyrus (Fig. 2B).

**Fig 2.**
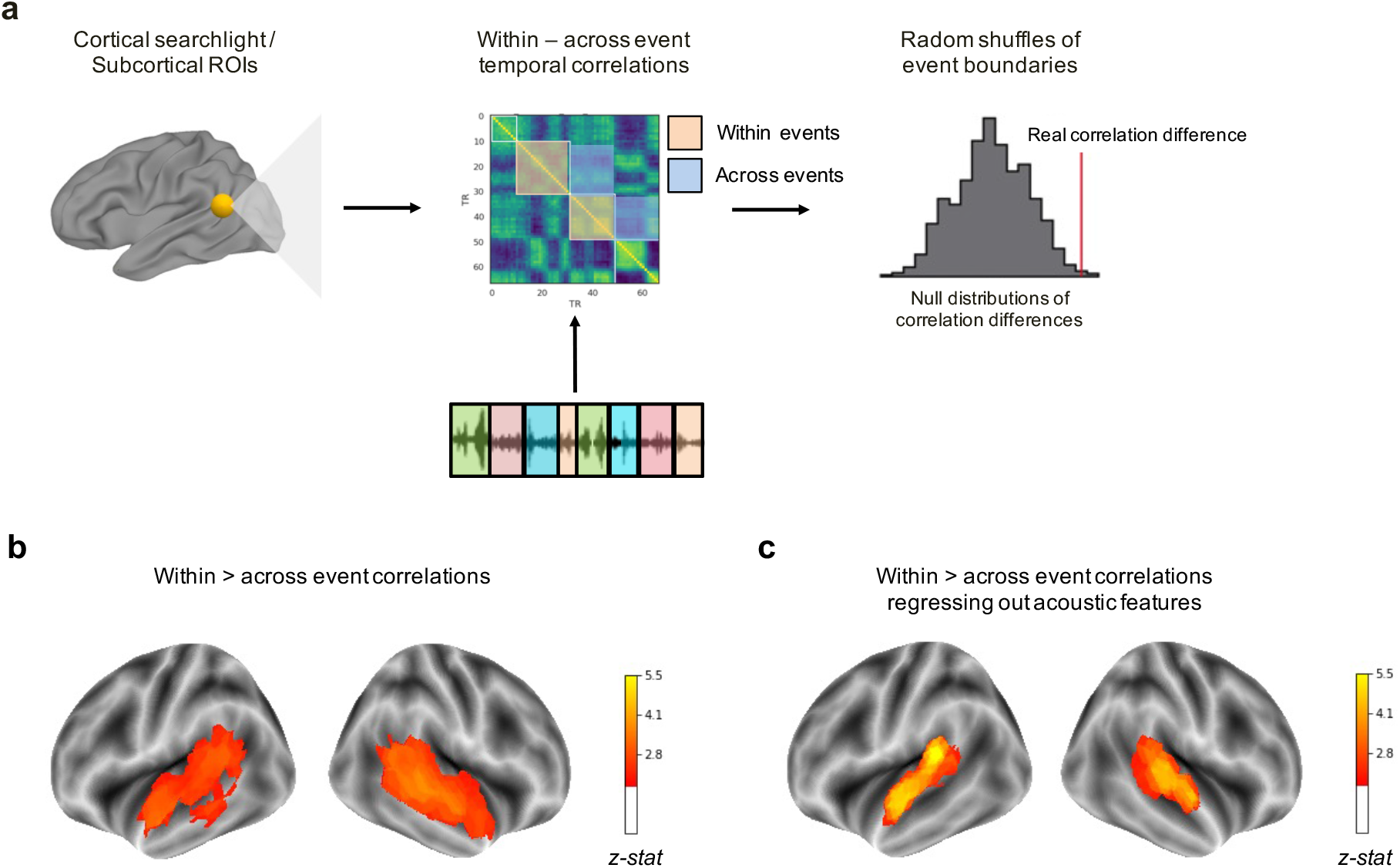
Brain regions sensitive to emotion transitions. A. Schematic of hypothesis-driven analysis for brain regions sensitive to emotion transitions by comparing within event vs. across event temporal correlations. B. Brain regions in which timepoint-by-timepoint correlations were significantly greater within a composer-defined emotional event as compared to across emotional events. C. Brain regions that show shifts at emotion transitions after acoustic features are regressed out. Colors correspond to z-scores across the cortical surface, relative to a null distribution (cluster-corrected p value < 0.05).

### Data-driven approach: HMM brain state transition results

HMMs can model sequences in brain patterns by assuming that states are characterized by periods in which brain activity patterns remain stable over time. HMMs have been used previously with neuroimaging data to show that our brains naturally transition between several different mental states that are similar across people, are spatially distinct, and are temporally ordered^19^. Here, we use this approach to uncover brain states that reflect emotion states induced by the music.

For each searchlight/ROI, an HMM-based event segmentation model was applied to brain data averaged across all participants. The number of events imputed into each HMM was set to the number of emotional transitions defined by the composers (16 for each piece). After fitting the HMM, we calculated the state entropy for each TR, which tells us the degree of certainty in the model of a transition at that moment in time. Finally, we averaged all entropy values that corresponded to the composer-defined musical transition timepoints and determined if these values were greater than what would have been expected by chance through permutation testing.

We found that likelihood of a brain-state transition was significantly greater than chance at composer-defined transitions in the bilateral auditory cortex, including the superior temporal and middle temporal gyrus, extending posteriorly into the left supramarginal gyrus and angular gyrus, and dorsally into the inferior parietal lobule and TPJ (Fig. 3B).

**Fig 3.**
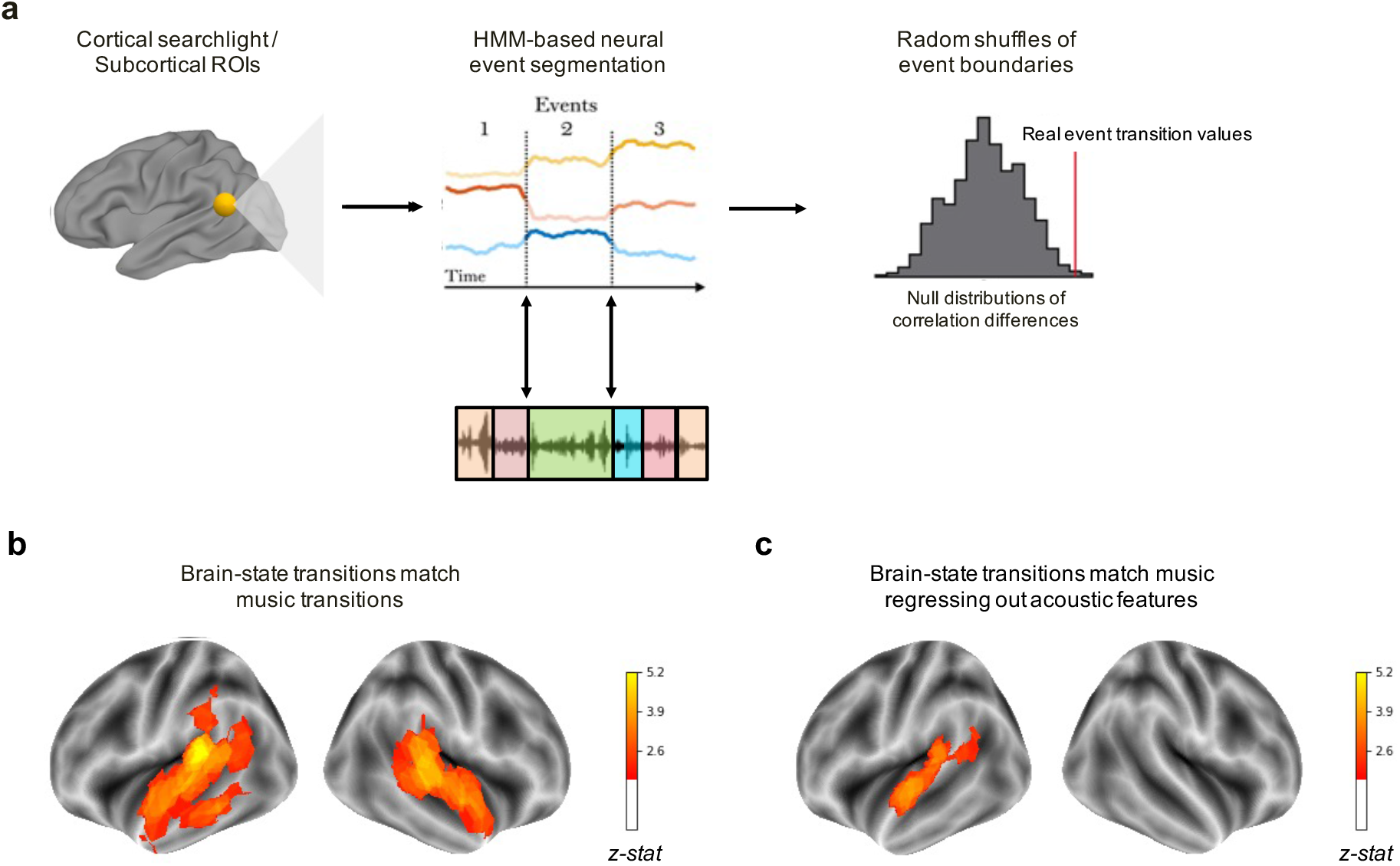
Brain-state changes driven by emotion transitions. A. Schematic of data-driven analysis for brain regions sensitive to emotion transitions using HMMs. B. Brain regions in which HMM-defined brain state transitions showed a significant match with composer-defined emotion transitions. C. Brain regions that are sensitive to emotion transitions after acoustic features are regressed out. Colors correspond to z-scores across the cortical surface, relative to a null distribution (cluster-corrected p value < 0.05).

The right hippocampus also showed higher entropy at composer-defined musical boundaries, though this effect did not survive correction for multiple comparisons across the 11 subcortical ROIs (*z-stat* = 2.39, p_uncorr_ = 0.008).

### Results when regressing out acoustic features

The two analyses presented above were repeated on data with acoustic features of the music pieces regressed out of the SRM feature space data first using the residuals from a linear regression model. These musical features are the same as those used in the encoding models and include information related to dynamics (rms), articulation (attack log), timbre (chroma centroid) and harmony. The hypothesis-driven largely mirrored the original findings and varied only in that the extent of the significant results in the temporoparietal cortex (see Fig 2C). Specifically, the right temporal pole and bilateral middle temporal gyrus no longer showed greater within-event vs across event temporal correlations. Furthermore, matches between HMM-defined brain state transitions and composer-defined emotion transitions were no longer significant in any part of the right hemisphere axis after acoustic features were regressed out of brain signal and were no longer significant in the left temporal pole and middle temporal gyrus (see Fig 3C).

### Do emotional features significantly predict time-varying activation patterns over and above acoustic features of the music?

To answer this question we constructed an encoding model, an approach in which stimulus features are used to predict BOLD signals. Measuring the relative performance of models that use different kinds of information about the stimulus can provide insight into the specific stimulus dimensions that drive neural activity in a brain region^23,24^. Here, we used this approach to determine if adding information about emotion ratings provides better prediction of BOLD signals, compared to a baseline model that uses only acoustic features of the music.

After scanning, participants listened to short excerpts extracted from each of the emotional events they heard during scanning and were asked to provide retrospective subjective ratings of how they felt the first time they heard to this moment within the piece. Specifically, they rated how happy, sad, anxious, calm, nostalgic, and surprised they felt as well as how much they enjoyed and how vividly they remembered this moment. Two separate regularized (ridge) regression models were employed to predict brain signal within a searchlight/ROI for each participant. One model included regressors for event-level musical/acoustic features, i.e. changes in tempo, dynamics, modality, timbre, and articulation across events. Another model included these same musical/acoustic features plus the 8 subject-specific emotional ratings collected post-scanning. We then identified regions that showed significantly greater correlations between predicted and actual signal in the full model as compared to the model with acoustic/musical features only.

We found that the full model with emotion features better predicted signal during music listening in vertices within the left ventromedial prefrontal cortex, left postcentral gyrus, and left visual cortex (Fig. 4).

**Fig 4.**
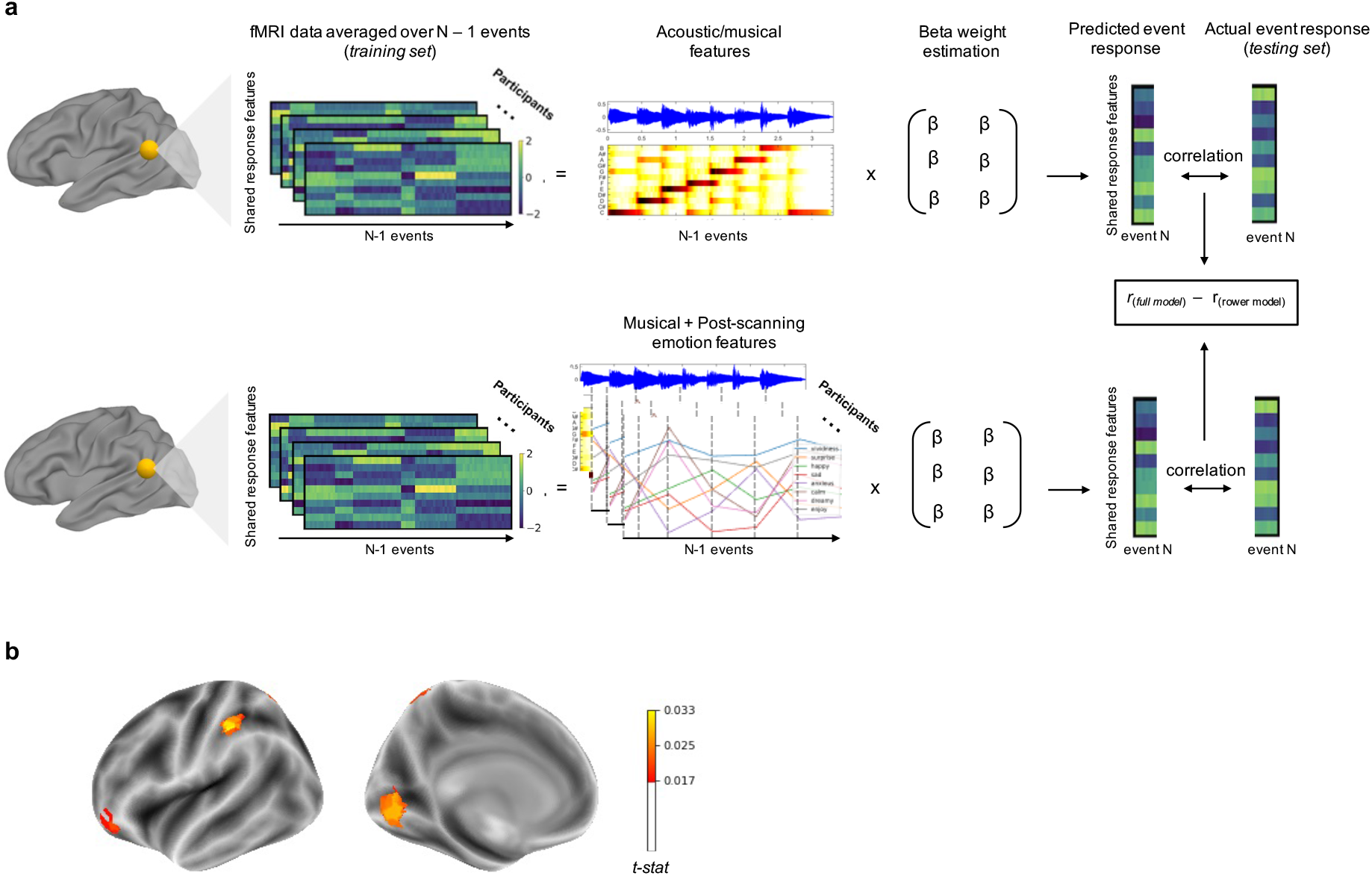
Brain activation patterns predicted by emotion vs. acoustic features. A) Schematic of the two encoding models and their comparison. B) Brain regions in which an encoding model with acoustic + emotion features showed greater correlation between predicted and actual brain signal as compared to the encoding model that contained only acoustic features. Colors correspond to differences in average r-values between the full (acoustic + emotion) model and the lower model (acoustic features only), and were cluster-size corrected at p < 0.05.

### How do brain states and their transitions vary as a function of the recent emotional context?

#### Systematic changes in behavioral ratings based on emotional context

We first assessed the degree to which emotion ratings of the music from the behavioral study varied as a function of condition, i.e. does how we report our emotions to a particular piece of music vary based on emotion that came before it. Specifically, for each music clip, we calculated the correlations between the post-behavioral test retrospective ratings for pairs of participants who heard that clip within the same condition (both version 1) as well as across conditions (one heard it in version 1 and another in version 2) and calculated differences in the mean of the within-condition pairwise correlations and the across-condition pairwise correlations. Across all clips, the within-condition correlation was significantly greater than the across-condition correlations (M_within_ = 0.303, M _across_ = 0.265, z-stat of difference = 2.0, p-value = 0.042), suggesting that subjective multivariate emotion ratings were systematically influenced by the prior emotional state. When averaging across emotional label, the calm and sad clips varied the most by context (Within *r_happy_* = 0.37, Across *r_happy_* = 0.35; Within *r_sad_* = 0.20, Across *r_sad_* = 0.15; Within *r_calm_* = 0.19, Across *r_calm_* = 0.15; Within 0.50, Across *r_anxious_* = 0.48).

In addition to the overall ratings of each event, we tested if the context manipulation influenced the time it took for participants to recognize and report feeling the intended emotion of each clip. For this analysis, we determined the time-to-peak for each event, operationalized as the moment when the number of participants who turned on the intended emotion for this event reached 90% of its maximum. The time to peak for positive events that were preceded by positive emotions was 9s faster than positive events preceded by negative events, which was determined to be statistically significant (*z-stat* = 1.99, *p-value* = 0.02). Negative events preceded by negative events were on average 2.3s faster to reach a peak in ratings than negative events preceded by positive events, a difference that was not statistically significant (*z-stat* = 0.49, *p-value* = 0.31).

#### Systematic changes in spatial brain patterns of emotional events based on emotional context

For each searchlight/ROI, we computed the average response pattern within each of the 35 events for each participant, and then calculated the correlations between participants for each event. We again binned these correlations based on whether the pair of participants heard that particular event within the same condition (both in piece A version 1, for example) or in different contexts (one in piece A version 1 and one in piece A version 2). We next tested which brain regions show significantly greater same-context pairwise correlations as compared to different-context pairwise correlations.

Significant differences in the spatial patterns averaged over emotional events that varied in the context were found bilaterally in the temporal lobe, including the primary and secondary auditory cortex (superior temporal gyrus) as well as the right anterior temporal lobe. Systematic changes were also shown in the right precentral gyrus and sulcus (Fig. 5).

**Fig 5.**
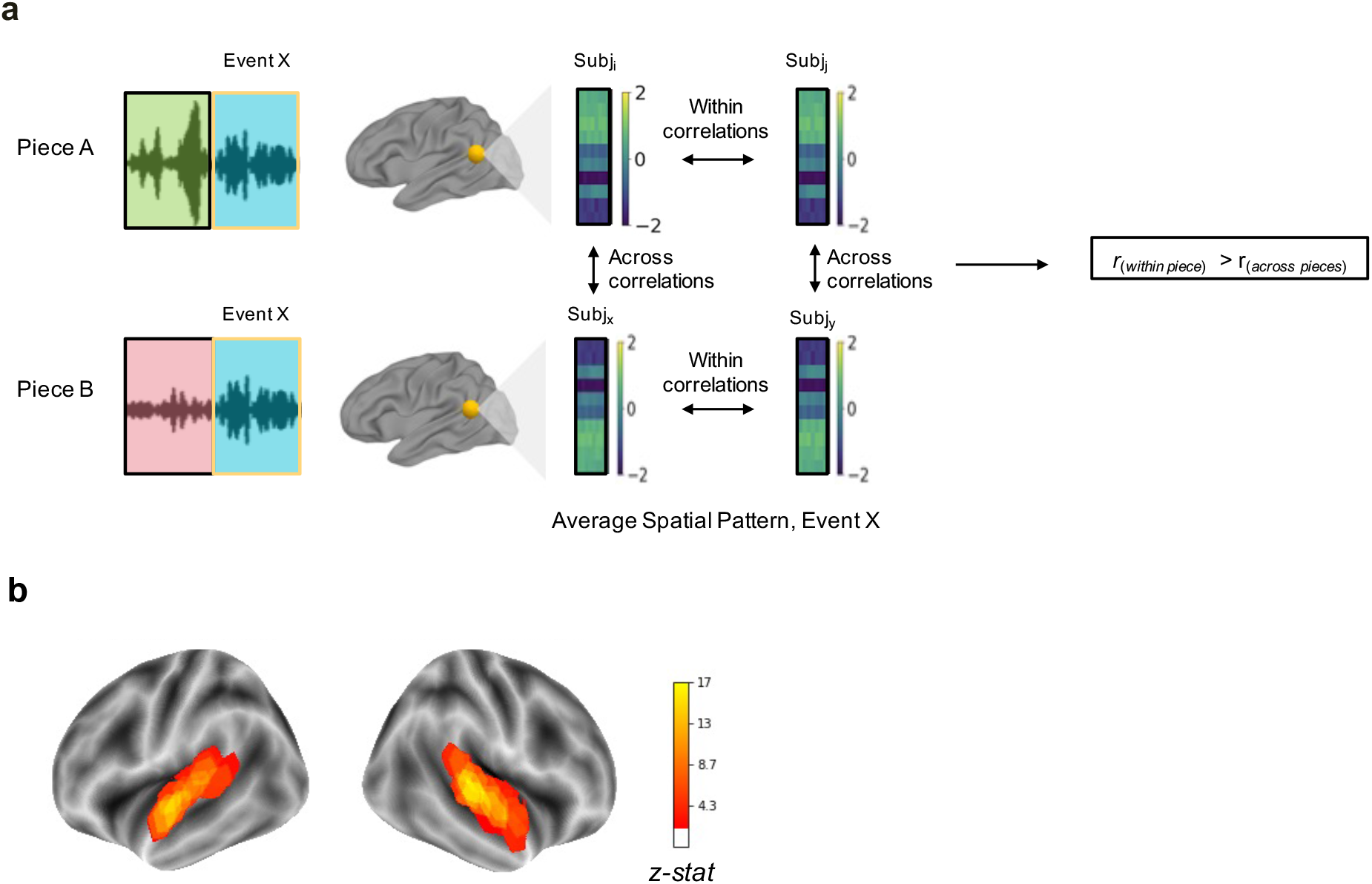
Context-based changes in spatial brain patterns. A) Schematic of pattern similarity analysis used to identify regions that show systematic differences in representation of emotional events based on condition (heard in the same piece vs. across pieces). B) Brain regions in which spatial patterns were significantly more similar in pairs of participants who heard the emotional event in the same condition (all in piece A or B, within group) as compared to pairs of participants who heard the music in different conditions (one in piece A other in B, across group). Colors correspond to z-stats of the ratio of across group vs. within group correlations as compared to a null model in which group membership was randomly permuted. Resulting statistical maps were cluster-corrected at p value < 0.05.

To avoid the possibility that differences were an artifact of fMRI signal spilling over from the event before (arbitrarily resulting in pairs of participants who heard the piece in the same context appearing to have activation patterns more similar due to the signal coming from the previous event), the analysis was run using only data averaged across the second half of each event; that is, not including any data that was temporally close to an emotion boundary/transition. The results were largely the same (see Fig. S2), suggesting that brain representation of emotions in the auditory cortex are sensitive to the emotional history in which the stimulus is encountered.

#### Systematic shifts in timing of emotion transitions based on emotional context

A measure of the “speed” of the transition from one emotional event to the next was calculated from the output of HMM models, fit to group-average data within a particular searchlight/ROI for two sequential emotional events at a time ^30^. We then calculated the difference in event transition timing for transitions with a valence shift(negative to positive or positive to negative) vs. emotional transitions within the same valence (positive to positive or negative to negative).

Significantly earlier (arriving to the next state earlier) transitions from same-valence contexts as compared to different-valence contexts were found in surface vertices corresponding to the right auditory cortex (including the superior temporal gyrus) and the left superior frontal gyrus (Fig. 6).

**Fig 6.**
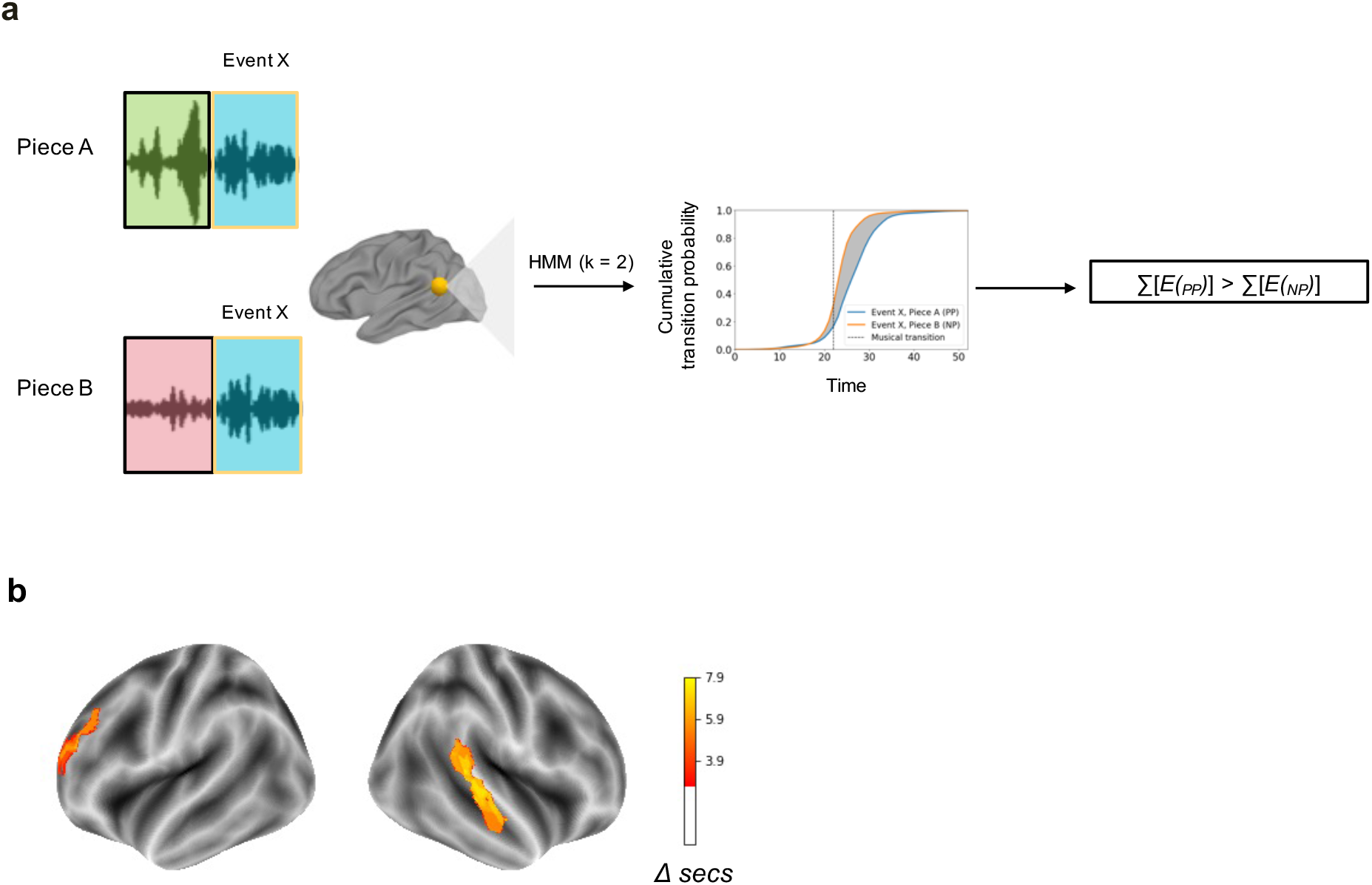
Context-based changes in temporal brain patterns. A) Schematic of temporal analysis with HMMs to identify how changes to the preceding emotional state altered the speed of the transition to the current state, separately for positive valence and negative emotional events. B) Brain regions in which HMM-defined transitions were significanty earlier when the event was preceded by the same valence as compared to a different valence. Colors correspond to the difference in seconds between same valence emotion transitions as compared to different valence transitions (same > different greater than), thresholded based on one-sample t-tests and cluster-size at p value < 0.05.

## Discussion

Emotions are dynamic by nature, reacting to our ever-changing environment in order to motivate adaptive behaviors. In order to more rigorously assess how our brains process emotional fluctuations and transitions, we developed novel musical stimuli that reliably induced various emotional states without visual and linguistic information that could confound with emotional information. We then tested if spatial and temporal brain patterns reflect emotion transitions evoked by music as well as how these transitions are altered by the emotional context in which the music is heard. We found evidence that several regions in the temporal and parietal lobes of the brain, including primarily sensory regions like the auditory cortex, show shifts in brain activation patterns that correspond to transitions between music-evoked emotional states. Furthermore, these patterns were systematically modulated by what emotional state preceded it. These results extend the role of the temporal-parietal axis as not only processing long-term narrative^31^ and musical structure^20,21^ as well as discrete musical-evoked^32^ and vocal emotions^33^, but high-level, contextually-dependent information regarding multidimensional emotion dynamics and their transitions.

Using a hypothesis-driven approach, we found that brain state transitions in voxels in the temporal lobe aligned with composer-defined emotion transitions. In combination with a data-driven analysis, we determined that emotion dynamics not only contribute to brain-state patterns in these regions, but are one of the primary drivers of brain responses to music. Previous research has shown that the temporal lobe plays an active role in both emotion perception and induction, in addition to processing of lower level perceptual aspects of sound^32^. Emotional labels associated with both musical and vocal sounds could be reliably decoded from voxels with the primary and secondary auditory cortex, suggesting that these regions represent the emotional content of sounds, independent of their specific acoustic properties^33^. Here, we extend these findings by showing that multivariate signals within the auditory cortex fluctuate between different stable states and that these state transitions are more likely to occur at moments when our emotional states also change. Importantly, Importantly, when acoustic features extracted from the music were regressed out of brain signal, the auditory cortex and superior temporal gyrus, except for the right auditory cortex in the case of data-driven results, continued to represent emotional changes in the music, suggesting that, at least in the left hemisphere, acoustic changes were not the soul driver of the time-varying patterns associated with emotional transitions.

Previous research has also highlighted the role of the temporoparietal axis, including the angular gyrus and TPJ, in representing temporal structure of music^20,21^ and narratives^31^, with short to long term information being represented hierarchically along the axis. However, it is unclear from these previous studies the nature of the temporal information that is being tracked. Our results argue that emotional changes are an essential signal for the brain when segmenting our continuous experiences. The superior temporal sulcus in particular has been theorized to track socially and emotionally relevant information over time ^31,34,35^. Here, we provide evidence for this theory by showing that emotional structure is one of the main organizing principles by which the superior temporal sulcus parses longer temporal experience.

To determine the degree to which changes in subjective emotional experiences is the main driver of fluctuations in the brain state patterns described above, we additionally ran encoding models to predict brain signals during music listening from a set of both subjective and musical features. While acoustic and musical changes in the music could reliably predict auditory signal, a model that included emotion features as well performed no better in auditory cortex, suggesting that the time-varying patterns that are sensitive to the composer-defined emotion transitions are most related to musical and acoustic changes in the music. Given the known role of temporal lobe regions for processing acoustical aspects of music^36^ and that it is not possible to completely dissociate musical changes from emotional changes, it is not too surprising that subjective emotional features add little to the model. However, the fact that several regions in the temporal lobe still reflect emotion transitions after acoustic data is regressed out from the signal suggests that the emotional content is still an important element in driving brain state changes in these regions.

Interestly, regions in which encoding models that included subjective feelings outperformed a more basic model with only acoustic features included left ventromedial prefrontal cortex, left postcentral gyrus, and visual cortex. The role of the ventromedial prefrontal cortex in affective experiences has been well documented^37,38^ and recently it was shown that brain states in the VMPFC align across participants during emotionally salient scenes of a popular TV show^12^. These results suggest that the VMPC has a very active role conferring our ongoing external experiences with affective meaning. This is in line with additional research showing that functional connectivity between the VMPFC and other brain areas changes during music-listening that is particularly emotional^39^ or particularly rewarding^40^, suggesting its role as a modulator/computational hub/gate-keeper for coordinating emotion responses and with perception of music.

The involvement of the postcentral gyrus and visual cortex in processing emotional changes to music is more surprising. The fact that subjective emotion features were better predictors of V1 activity could be due to mental imagery involved in music-listening, which enhanced

the emotional experience. This hypothesis seems more likely when considering that the composers who wrote our musical stimuli were all trained in film scoring, which is designed to accompany and enhance visual images. Previous studies have shown increased V1 activity and/or connectivity during music listening that is particularly emotional^39,41^ as well as in listeners that are more empathic, and therefore more likely to experience strong emotions to music^39,41^. It may well be that experiencing vivid visual images during music listening allows for stronger empathic connections to the composer, characters, or events of the music, which, in turn, fosters more powerful emotional responses. Given that the full feature encoding model included self-report ratings of vividness as well as emotional intensity, and that post-scanning ratings of vividness were correlated with ratings of emotional intensity, it could be that dynamic patterns in the visual cortex that our model is able to predict reflect more vivid mental imagery that coincide with more intense feelings.

Not only did patterns of brain activity along the temporal parietal axis represent emotion transitions, spatial patterns during emotional events within these regions were sensitive to the emotional context in which the music was heard. Specifically, spatial correlations (averaged across emotional event lengths) in the temporal lobe were significantly more similar in pairs of participants who heard that emotional event with the same prior context as compared to pairs of participants who heard that emotional event with different prior contexts. This suggests that, contrary to previous research limiting its role to the processing of local, lower-level, shorter-timescale features of musical structure, the auditory cortex is sensitive to the previously-established context in which music is heard. While it is possible that this increased similarity is an artifact as a result of fMRI-signal spilling over from the event before (which would make pairs of participants who heard the piece in the same context look more similar because the previous event was also the same), we re-ran the analysis using only data averaged across the second half of each event, that is, not including any data that was temporally close to an emotion boundary/transition. The results in the temporal lobe were largely the same, suggesting that the representation of musical-evoked emotions in the auditory cortex is sensitive to the context in which the music is encountered. Future research will be needed to determine the exact timescale on which preceding context influences auditory representations of music.

Interestingly, in addition to this change in representation, the right superior temporal lobe also showed a temporal effect of context. Specifically, the timing of brain-state transitions between emotional events in these regions varied as a function of the valence of the preceding event. Behavioral evidence indicated that the time until which most participants agreed that a musical event made them feel a particular emotion varied depending on what type of musical emotion came before. Specifically, people were more likely to feel positive emotions when the preceding event was also positive (joyful and calm vs. sad/anxious). This pattern was also reflected in the timing of HMM-defined brain state transitions in the auditory cortex, since the model detected transitions earlier in time when the valence of the preceding event was the same as the current event. While previous studies using similar approaches have shown how repeated viewing of a movie^30^ and aging^28^ can temporarily shift activity patterns in a way that reflects changes to our subjective experiences, this is the first study, to our knowledge, to show that changes in our subjective experience of a piece of music can alter brain representations of the associated emotion.

Despite our initial hypothesis, subcortical areas of the brain that have been reliably shown to respond during emotion processing (e.g. amygdala, thalamus, hypothalamus, caudate, striatum) did not show time-varying brain patterns that reflected music-evoked emotion dynamics. While changes in these regions appear to be reliably predicted by emotional responses evoked by short videos^23^ and in some studies that used music designed to induce emotions^32,42^, other recent evidence found that not all limbic regions reliably represent music-induced emotions^43^ or were activated as strongly as by other rewarding stimuli ^44^. Furthermore, the majority of studies finding significant activation of subcortical regions corresponding to music-evoked emotions averaged signal over the duration of the piece and compared this average signal to some control condition (scrambled music, sine tones, silence, or music designed to convey another emotion). It is therefore possible that increased signal in these regions is a reflection of an overall affective response, but that they do not show the types of patterns that our analyses are designed to pick up in these analyses, i.e. stable patterns within an emotional event followed by rapid shifting to a new stable patterns. Further work will be needed to determine whether dynamic emotional experiences evoke different kinds of dynamics in subcortical areas, such as transitory responses at boundaries or ramping activity throughout events^17,45^.

This study has several limitations that are important to address. First of all, due to the limitations of statistical power, we could not assess whether emotional changes of a certain type (a positive valence emotion to a negative valence emotion, or a high arousal emotion to a low arousal emotion) were the main drivers of brain transitions. Previous research has suggested that changes in context elicit changes in arousal, which segment our memories into separable events^46^. It could be that specific “types” of emotion transitions (e.g. negative to positive state) or only more salient/arousing transitions are driving the event patterns in different ways, though given the limited number of each type in our stimulus, it was not possible to test the quality of the event transition that may be driving the brain-state changes. Finally, most of the analyses presented here focus on group-level statistics, though it is possible that emotion dynamics are not stable or consistent *across* participants. Capturing these individual differences is beyond the scope of this paper, and could account for why traditional emotion regions did not emerge in these analyses. Follow-up investigations will try to assess individual differences in emotional experiences to music using within-subject analyses and richer self-report measures.

In sum, using novel music composed specifically for the purposes of driving listeners through different emotional states in an optimized period of time, we show that regions along the temporoparietal axis show spatiotemporal patterns that reflect the changing emotional experiences to music. Specifically, we found stable brain patterns within the primary auditory cortex, superior temporal gyrus and sulcus during an emotional event that rapidly shifted to a new stable pattern during emotion transition periods in the music. Activation in these regions also showed altered spatial and temporal patterns to the same pieces of music that were heard in different emotional contexts. The findings suggest a role of the temporoparietal axis in integrating changing acoustic input with our changing internal states, highlighting a potential mechanism by which our emotions fluctuate in everyday life and treatment target for when such fluctuations go awry in the case of mental illness.

## Methods

### Ethics information

This study was conducted under an approved study protocol reviewed by the Columbia University Institutional Review Board (IRB AAAS0252). Informed consent was obtained from all human participants. Participants received monetary compensation ($20/hour) for their time.

### Stimuli development and behavioral validation

To develop non-linguistic emotional stimuli suitable for fMRI, we hired three film score composers, all graduate students at New York University’s Film Composition program, to write two original pieces of music. The composers were asked to divide the pieces into sections, where each section conveyed a single emotional category. Emotional categories were selected based on dimensions identified from a large, cross-cultural musical corpus in which people reported both dimensional and categorical emotional responses^47^. After discussion with the composers regarding what was feasible given the musical constraints, we arrived at five distinct emotional categories: sad/depressing, anxious/tense, calm/relaxing, joyous/cheerful, and dreamy/nostalgic. We also asked that each emotional category be revisited 7 times across the two pieces, using different musical elements during each recapitulation. The length of each section and the timing of the transition from one emotion to another we left up to the composer, though we asked that each section be no less than 30s. Additionally, the tempo was set to be the same as (or a multiple of) the fMRI pulse sequence (80/160 BPMs) and composers were instructed to use up to four separate voicings/instruments (violin, piano, guitar, cello).

Importantly, the composers were able to write these musical events modularly, so that the ordering could be shuffled without disturbing the natural flow of the overall piece. This allowed us to manipulate emotional context in a systematic way. Specifically, 32 events were divided into two distinct musical pieces (A and B), each with 16 unique musical events (4 emotions x 3-4 examples, ~15min in length). Musical interludes (4-12s) were written by the composers and inserted in between event transitions to musically link one event to another and to allow the piece to sound like a cohesive whole when played continuously. While the ordering of the specific events was left up to the composers, the pieces were constructed to ensure that the number of events preceded by an exemplar of the *same* valence (joyful and calm considered positive and sad and anxious considered negative) was equal to the number of events preceded by an exemplar of a *contrasting* valence (see Fig. 1A). Given that nostalgia is considered a mixed emotional state with both positive and negative aspects ^48^, we did not have specific hypotheses about its effect on subsequent emotional events and therefore only included 4 of the dreamy/nostalgia clips, one at the beginning and end of each piece. This resulted in 14 emotion transitions of the same valence (positive to positive or negative to negative) and 14 emotion transitions of contrasting valence (positive to negative or negative to positive) across the two pieces (7 of each type in each piece). From this initial set of two pieces, the composers created an alternative version of each, using the same events, but re-ordered them so that any event that was previously preceded by a contrasting valence emotion was now preceded by a same valence emotion, and vice versa. For a list of all events and their lengths, see the supplementary materials. All audio files are made available to reviewers on OSF (https://osf.io/a57wu/?view_only=996150c5d6fc49c1a004c78ef2f852d3).

Several additional audio engineering changes were made to the stimuli to increase their suitability for MRI. The frequency of the repeating sounds of our particular SEIMENS MRI machine corresponded to a C note. Therefore, all of the music was transposed to the key of C, a pitch shift of a whole step down from where it was original written, ie. the key of D. Additional increases of the gain of certain bandwidths were also adjusted to allow for a more optimal listening experience with the headphones inside MRI.

### Assessing the validity of emotional states and their transitions with independent ratings

In order to validate the emotion transition timepoints and the emotions they were intended to induce, subjective emotion measures were collected via a custom open source web application built using JavaScript (http://www.jonaskaplan.com/cinemotion/) from an independent group of participants. The tool instructed participants to listen to a piece of music and to think about what emotion they are feeling in response (not how they think the performer/composer is feeling). While the music is playing, participants were instructed to select one of five possible (“happy”, “sad’, “anxious”, “nostalgic”, and “calm”) buttons to “turn on” that emotional label when they felt it and to press it again to “turn off” that emotional label when they no longer felt that particular emotion. The label, onset time, and offset time were recorded continuously. Each participant listened to only one of the 4 possible 15-minute pieces, divided into three ~5-minute sections with a self-paced break in between each to maintain focus. The breaks occurred within the middle of an emotional event, not at transition points. In addition, at the end of the final section, the ending of the piece transitioned into a completely different piece (*Blue Monk* by Thelonious Monk). This contrast was used as an attention check: any participant that did not turn off or on any of the 5 buttons within a window of time (1.5 before to 5.7s after) around the transition to this new piece was removed from the analysis.

To determine the number of people needed to rate each piece, we used results from a previous dataset of N = 80 people rating music clips designed to induce happiness or sadness^11^. With 100 bootstrapped samples of N participants randomly (ranging from 10-80) sampled from this dataset, the average RMSE between mean ratings with 35 people and the overall mean ratings with 80 people explained 67% of the variance, compared to 43% with 25 and 79% with 45. This suggests that ratings from 35 people is sufficient to have a reliable estimate of how people report feeling in response to emotional music and provides a reasonable trade-off between cost/time and variance explained.

Based on this analysis, we recruited 40 participants to listen to each of the four pieces with the assumption assuming that ~10% of subjects would need to be excluded. After removing participants that did not press any buttons for the duration of the piece or failed the attention check, we analyzed the ratings from 36 people who heard piece A version 1, 36 people who heard piece A version 2, 35 people who heard piece B version 1, and 35 people who heard piece B version 2.

At the moments of transitions between emotional events, as identified by the composer, we counted the number of raters that turned on or off any emotion at every 1s timepoint. We then counted the number of ON/OFF selections at the transition intervals. To determine if people were more likely to press the emotion buttons during the transition periods than not, we randomly shuffled the order of the events 1000 times and for each permutation, we recalculated the number of people who reported an onset of the equivalent emotion at the randomly permuted onset time to create a null distribution. Across all four pieces, the number of raters who turned ON/OFF an emotion at the transition points was significantly greater than at random timepoints (Piece A1 *z-stat:* 3.6, p-value = 0.0002; Piece A2 *z-stat:* 3.21, p-value = 0.0006; Piece B1 *z-stat:* 4.32, p-value < 0.0001; Piece B2 *z-stat:* 4.15, p-value < 0.0001). Furthermore, for every transition point throughout all 4 pieces, there were more raters identifying changes in the 3s after the transition than in 95% of permuted transitions.

We next calculated the average number of people that had selected the composer-intended emotion during all the timepoints within each emotional event. We computed a null distribution by randomly shuffling the emotional event labels 1000 times and recalculating the mean. The average number of people that reported experiencing that emotion throughout the duration of the event was significantly greater than chance for all emotional categories, except for “nostalgia” (calm: mean = 17.63, *z-stat* = 2.28, p-value = 0.01; happy: mean = 17.22, *z-stat* = 4.40, p-value < 0.001; sad: mean = 16.79, *z-stat* = 3.38, p-value < 0.001; anxious: mean = 19.16, *z-stat* = 5.53, p-value < 0.01; nostalgic: mean = 11.16, *z-stat* = 1.09, p-value = 0.14).

### Post-behavior retrospective recall task

After listening and rating to the ~15-minute piece of music, participants completed an online survey on Qualtrics that utilized retrospective behavioral sampling^49^. Specifically, for each emotional event in the full piece of music, we extracted three unique 10s excerpts taken from the beginning, middle and end of the emotional event (see below for more details).

Participants then listened to one of these three clips (exactly one from each event they heard during scanning, i.e. 32 events) selected randomly and presented in a random order. After listening to the clip, they were then asked to focus their memory on the first time they heard that particular moment in the music during scanning, including no more than a few moments before and after it and to rate 1) how *vividly* they remember this moment in the piece on a 7-point likert scale. If they do remember that particular moment (ratings > 1), they were subsequently asked to rate 2) how *surprising/unexpected* that moment in the music was the first time they heard it, 3) how *happy/joyous* did that moment make them feel; 4) how *sad* did that moment make them feel, 5) how *anxious/tense* did that moment make them feel, 6) how *calm/relaxed* did that moment make them feel, 7) how *dreamy/nostalgic* did that moment make them feel, and 8) how much did they *enjoy* this moment of the piece. Participants listened to exactly one 10s clip from each of the emotional events from the piece that they heard, as well as one additional clip from the piece they did not hear as an attention check, for a total of 17 clips.

The clips were created using pyDub package, a Python-based library used for audio manipulation (https://github.com/jiaaro/pydub). Each emotional event was segmented into three 10s clips, evenly spaced throughout the duration of the event. To avoid the impact of onset/transition spillover, the first 3s and last 2s of the emotional event were not included. Because emotional events varied in length, this meant that the gap between successive clips varied and was determined by taking the entire duration of the event and finding the gap length that evenly spaced the clips throughout the duration of the event. During the survey, participants only heard 1 of the 3 possible clips from each event, randomly chosen and balanced across participants.

### fMRI task, acquisition, and preprocessing

During scanning, participants listened to two full-length pieces of music (A and B) with no explicit instructions other than to listen attentively and restrict movement as much as possible (see Fig. 1B). Which version of piece A and B, as well as the order of presentation of the two stimuli, was counterbalanced across participants. In between the two music-listening sessions, participants watched a ~12.5 minute audio-visual movie (*Rhapsody in Blue* from *Fantasia 2000*), which was used for functional alignment^29^.

MRI images were acquired on a 3T Siemens Prisma scanner using a 64-channel head coil. T2*-weighted echoplanar (EPI) volumes were collected with the following sequence parameters: TR = 1500 ms; TE = 30 ms; flip angle (FA) = 90°; array = 64 × 64; 34 slices; effective voxel resolution = 2.5 × 2.5 × 2.5 mm; FOV = 192 mm). A high-resolution T1-weighted MPRAGE image was acquired for registration purposes (TR = 2170 ms, TE = 4.33 ms, FA = 7°, array = 256 × 256, 160 slices, voxel resolution = 1 mm^3^, FOV = 256). Each of the two music-listening scans consisted of 607 images (6s of silence before the music begins, 896s/597 images of music listening, followed by 9s of silence at the end). The movie-watching scan was acquired with identical sequence parameters to the EPI scans described above, except that the scans consisted of 496 images (744s).

MRI data was converted to Brain Imaging Data Structure (BIDS) format using in-house scripts and verified using the BIDS validator: http://bids-standard.github.io/bids-validator/. The quality of each participant’s MRI data was assessed using an automated quality control tool (MRIQC v0.10)^50^. MRIQC creates a report for each individual scan based on assessment of movement parameters, coregistration, and temporal signal-to-noise (tSNR) calculations. We visually inspected the assessment reports for each participant to ensure adequate coregistration and fieldmap correction.

Functional data included in this manuscript come from preprocessing performed using FMRIPREP version 20.2.1^51,37,38^ [RRID:SCR_016216], a Nipype^52^ [RRID:SCR_002502] based tool. Each T1w (T1-weighted) volume was corrected for INU (intensity non-uniformity) using N4BiasFieldCorrection v2.1.0^53^ and skull-stripped using antsBrainExtraction.sh v2.1.0 (using the OASIS template). Brain surfaces were reconstructed using recon-all from FreeSurfer v6.0.1^54^ [RRID:SCR_001847], and the brain mask estimated previously was refined with a custom variation of the method to reconcile ANTs-derived and FreeSurfer-derived segmentations of the cortical gray-matter of Mindboggle^55^ [RRID:SCR_002438]. Spatial normalization to the ICBM 152 Nonlinear Asymmetrical template version 2009c^56^ [RRID:SCR_008796] was performed through nonlinear registration with the antsRegistration tool of ANTs v2.1.0^56,57^ [RRID:SCR_004757], using brain-extracted versions of both T1w volume and template. Brain tissue segmentation of cerebrospinal fluid (CSF), white-matter (WM) and gray-matter (GM) was performed on the brain-extracted T1w using fast^58^ (FSL v5.0.9, RRID:SCR_002823).

Functional data was motion corrected using mcflirt (FSL v5.0.9)^59,60^. Distortion correction was performed using an implementation of the TOPUP technique^61^ using 3dQwarp (AFNI v16.2.07^62^). This was followed by co-registration to the corresponding T1w using boundary-based registration^59^ with nine degrees of freedom, using bbregister (FreeSurfer v6.0.1). Motion correcting transformations, field distortion correcting warp, BOLD-to-T1w transformation and T1w-to-template (MNI) warp were concatenated and applied in a single step using antsApplyTransforms (ANTs v2.1.0) using Lanczos interpolation.

Physiological noise regressors were extracted applying CompCor^63^. Principal components were estimated for the two CompCor variants: temporal (tCompCor) and anatomical (aCompCor). A mask to exclude signal with cortical origin was obtained by eroding the brain mask, ensuring it only contained subcortical structures. Six tCompCor components were then calculated including only the top 5% variable voxels within that subcortical mask. For aCompCor, six components were calculated within the intersection of the subcortical mask and the union of CSF and WM masks calculated in T1w space, after their projection to the native space of each functional run. Frame-wise displacement^64^ was calculated for each functional run using the implementation of Nipype.

Many internal operations of FMRIPREP use Nilearn^65^ [RRID:SCR_001362], principally within the BOLD-processing workflow. For more details of the pipeline see https://fmriprep.readthedocs.io/en/20.2.1/workflows.html.

Additional nuisance regressors were regressed out of the data, including six scan-to-scan motion parameters (x, y, z dimensions as well as roll, pitch, and yaw), their derivatives, CSF and WM signal, framewise displacement, and the first five the first five noise components estimated by aCompCor^39^. High pass temporal filtering (0.008 Hz) was applied using discrete cosine bases. The resulting whole-brain time series were then z-scored within subjects to zero mean and unit variance. All preprocessing steps were performed using custom scripts written in Python that incorporate a variety of packages, including *Brainiak, Nibabel, Nilearn, and Scikit-learn*.

### Functional alignment using the shared response model

Prior to any further analyses, to account for the fact that anatomical alignment techniques may be insufficient for aligning fine-grained spatial patterns across individuals, we used a shared response model (SRM) to functionally align regions into a common space^66^. Specifically, we fit the model using brain activation in response to an audio-visual movie without lyrics (*Rhapsody in Blue* by from the movie Fantasia 2000) and applied it to brain patterns recorded during the music-listening task. The model determines a linear mapping (from voxels to shared features) between an individual’s functional response and a shared response that is well-aligned across subjects^35^. Specifically, the weight matrix (features by voxels) fit to the movie data was used to transform raw voxel activity (voxels by time) during music listening into a shared feature space (features by time). To simplify subsequent analyses, the number of features was set to be consistent across all ROIs/searchlights, independent of the number of voxels within the ROIs. We chose to set the number of features to be 10% of the size of the largest ROI, yielding 80 features.

### Post-scanning retrospective recall task

After scanning, participants completed the same retrospective behavioral sampling survey on Qualtrics as the behavior participants (see above). For the excerpts that they remember hearing during scanning, participants provided ratings for how happy, sad, calm, anxious, surprised, and nostalgic they felt as well as how much they enjoyed that moment in the piece the first time they heard it. The ratings were used as input features for encoding models (see Fig. 1C).

### Inclusion/Exclusion Criteria

Data collection and analysis were not performed blind to the conditions of the experiments. That being said, all critical conditions are within-subject and within-run and the experimenter did not interact with the participant during scanning, except to ensure the safety of the participant. Participants were recruited through flyers posted throughout the Greater New York area and online. To qualify, interested individuals had to be between the age of 18-55, native English-speakers, mostly right-handed (as determined by the Edinburgh Handedness Questionnaire), with normal hearing. Additional exclusion criteria included: basic MRI contraindications (metallic implants, pacemaker, pregnancy), a history of psychosis, a history of electroconvulsive therapy, a history of brain disorders, including stroke, tumor, infection, epilepsy, degenerative diseases and head trauma, currently taking (or taken within the last month) medications that target the central nervous system (neuroleptics, anticonvulsants, antidepressants, benzodiazepines), and any diagnosis of learning disabilities.

Three participants were removed due to technical issues and one participant was excluded due to excessive movement (more than 20% of TRs for each session exceed a framewise displacement of 0.3 mm^67^. Finally, because our main analysis involved group-averaged data, we removed 4 functional runs in which participants showed particularly idiosyncratic brain data, operationalized as having an average pairwise ISC less than 2 standard deviations below the mean ISC across all participants.

### Which brain regions track transitions from one emotional state to another?

For all analyses, we used a multivoxel searchlight approach, in which data from circular groups of vertices on the cortical surface (radius 11 vertices/ ~15mm radius, with each vertex covered by 14 different searchlights) were iteratively selected for analysis^47^. We additionally ran the respective models/analyses on 11 subcortical regions of interest, including the left and right thalamus, striatum (caudate and putamen), pallidum, hippocampus, amygdala, and the bilateral nucleus accumbens, as defined by the Freesurfer subcortical parcellation. In each of the below analyses, to correct for multiple comparisons across searchlights, we performed a clustering threshold approach, for which we re-ran the given analysis 1000 times with null data, calculated the number of adjacent vertices that were statistical significant at the p = 0.05 uncorrected cutoff (to form clusters), and took the max cluster size for each permutation. We then determined the cluster sizes of our real data in the same way and determined how many of those were greater than 95% of the null clusters (cluster-threshold = 0.05).

### Hypothesis-driven approach

To identify brain regions sensitive to emotion transitions as defined by the composer, for each searchlight/ROI, the correlation between the patterns of shared features (from hyperalignment) were computed for all pairs of time points. We then took the average correlation between all TRs that were within an emotional event and all TRs that spanned an emotional event (i.e. between event t and an adjacent event t + 1). The statistical significance of this across-vs. within-boundary correlations was calculated by randomly shuffling the boundaries of events (preserving event lengths) and re-calculating the within vs. across boundary correlations for each region^68^. Cluster-thresholding was then applied to the resulting statistical map (see Fig. 2A).

### Data-driven approach

While the model-based approach affords us more statistical power, it does not tell us whether emotion transitions are the dominant factor driving pattern transitions in a brain region. We therefore supplemented the above findings with data-driven, generative models that try to learn latent “states” as well as their transitions based on the patterns of recorded brain activation. Combined with above, these results illuminate if primary brain pattern shifts are emotion-driven, not simply that emotional events contribute to pattern shifts. For this, we used an HMM-based event segmentation model^68^, which assumes that participants experience a sequence of discrete events while processing a naturalistic stimulus and each of these events has a discrete neural signature. The model also assumes that all states should be visited at least once and that all participants end in a particular state. For each time point, the model determines the likelihood that a region, based on signal across the shared feature dimensions, is in a particular state, assigning it a value between 0 and 1. Importantly, the model does not assume that events have the same length.

For each searchlight/ROI, the event segmentation model was applied to data in SRM space, averaged across all with the number of events set to the number of emotional transitions defined by the composers (16 for each piece). After fitting the HMM, we obtain an event by timepoint matrix for each piece, giving the probability that each timepoint belongs to each event. For each TR, we then take the entropy (using the scipy stats function) across the probability distribution, which tells us, for each TR, the likelihood of a boundary switch. We then calculated this entropy value at the moments of composer-defined transitions and determined if these values were greater than what would have been expected by chance through permutation testing. A null-distribution of entropy values was created by shuffling the timing of the behavioral events (preserving their lengths) 1000 times and re-calculating the entropy values at those new events. The average entropy at real event transitions was then compared to the average entropy value for null distribution to calculate a z-statistic and a subsequent p-value. Cluster-thresholding was then applied to the resulting statistical map (see Fig. 3A).

### Do emotional-categories significantly predict brain activation patterns above features that pertain to acoustics of the music or subjective experiences?

To evaluate the sensitivity of brain regions to changes in higher-level emotional dimensions (self-report emotional intensity, vividness, enjoyability, and surprise) as compared to lower-level musical and acoustic dimensions (instruments/timbre, dynamics, harmony/melody, tempo), regularized (ridge) regression was used to predict the time course of each feature from in shared space within a region. Ridge regression allows us to include multiple, highly-correlated predictors into a single model^69^. High-level emotion features included subject-specific ratings for the post-scanning recall questions (vividness, enjoyability, surprise, happy, sad, anxious, calm, nostalgic). Since each participant rated three different sections from each emotional event, the average of the three ratings within each subject was used. The low-level, musical features included the four features given from the composers: the time signature of each event (4/4 or 3/4), the BPMs (80 or 160), modality (major or minor), and the percentage of time within the event that each of the four instruments was heard (1 regressor for each instrument with values ranging from 0 to 1), as well as several acoustic features extracted for each event using the *librosa* Python package^70^: mean and std of root mean squares (RMS, dynamics), mean and std of the log of the attack phase of the envelope of the signal (articulation), the mean and the std of the per frame chroma centroid/chromogram center (pitch/melody), and the mean and std of harmonic change between consecutive frame (harmony, difference in harmonic content between consecutive frames)^71–73^.

For each subject, all regressors (brain, emotion, and musical) were z-scored, resulting in a feature by event matrix of vectors. For training and test, we used leave-one-out crossvalidation, in which one event was left out during each fold. For each fold, to find the optimal regularization parameter and avoid overfitting, an inner cross-validation was used on the training set, again leaving one event out for each fold and calculating the regularization parameter (20 possible alpha values geometrically spaced between 1 and 100000) that maximizes the correlation between actual value of the 80 shared response features for the held-out validation event and the predicted values (see Fig. 4A). This alpha value was subsequently used to predict share response feature values from the training feature set. Finally, we calculated correlation between the testing event brain data and the predicted values.

To determine the brain areas that were specifically sensitive to subjective emotional responses to the music, we ran two separate encoding models, one that included both emotion and musical features and one that included only musical features. We then tested which brain areas showed correlation values between the predicted and actual signal that was significantly greater in the full model (emotion + musical) as compared to the music only model. Statistical significance was computed using a one-tailed t-test to determine if the mean difference in correlations (full model minus musical model) was significantly greater than 0 across participants for each surface vertex. T-values were corrected using cluster thresholding (as described above).

### How do brain states and their transitions vary as a function of the recent emotional context?

#### Assessing the context manipulation on the time course of emotion ratings

To assess if the context influenced the patterns of emotional responses to the music, for each event that a participant heard, we constructed the matrix of the participant’s ON/OFF ratings (1/0) across all 5 emotional categories for all timepoints within the event. We then split participants into four groups: two groups (split-half) who heard the event in version 1 (V1_H1_ V1_H2_) of the piece and two groups (split-half) that heard the event in version 2 of the piece (V2_H1_ V2_H2_). For each musical clip, we next calculated the Pearson correlations between the ratings matrices for pairs of participants within the same condition (V1_H1_-V1_H2_ and V2_H1_-V2_H2_) as well as across conditions (V1_H1_-V2_H1_, V1_H1_-V2_H2_, V1_H2_-V2_H1_, and V1_H2_ - V2_H2_) and took the mean of the within-condition pairwise correlations and the across-condition pairwise correlations. Within vs. across condition pairwise correlations were compared to 1000 null correlations computed by shuffling the condition (clip version) distribution in which the participant condition was randomly shuffled 1000 times.

To determine if the context manipulation additionally influenced the time it took for participants to recognize and report feeling the intended emotion of each clip, we calculated the time-to-peak for each event, operationalized as the moment when the number of participants who turned on the intended emotion for this event reached 90% of its maximum. Only events in which at least 30% of the total participants turned on the intended emotion were included in this analysis, which excluded one calm clip. Across each emotion label, the average time-to-peak was fastest for anxious clips (M = 9.93s) followed by calm (M = 10.92s), happy (15.71s), and the slowest for sad (22.14s).

To statistically determine how the change in emotional context, i.e. the preceding emotion, modulated the time to peak, we first categorized all events across the two pieces into one of 4 conditions: a positive emotional event preceded by a negative emotional event (NP), positive emotion preceded by a positive emotion (PP), negative emotion preceded by a positive emotion (PN), and negative emotion preceded by a negative emotion (NN). This analysis excluded all nostalgic clips and the first events of each piece, resulting in 14 events in each of the 4 categories across all four pieces. We next created a null distribution by randomly permuting whether or not the event was preceded by a negative or positive emotion for each valence separately.

#### Systematic changes in spatial patterns of brain activation for emotional events based on context

To evaluate how preceding context influences brain representations of emotions, we assessed the degree to which brain activation pattern were systematically altered based on the emotional event that came before it. To this end, we quantitatively compare the pattern evoked by an event in all pairs of subjects. If context systematically alters brain patterns that correspond to a particular emotion, then subjects who experienced an event in the same context (both heard it in piece A, for instance) will have more similar patterns of activation. To test these hypotheses, within a particular ROI/searchlight, we first averaged the data across time points within each event, resulting in one pattern of SRM feature-wise activity per each emotional event. We did this for each participant individually and then calculated the across-subject pairwise Pearson correlation for each of the 35 events (see Fig. 5A). For each event, we then binned pairs of participants depending on which version of the two pieces they heard, that is, if the context in which that participant heard an emotional event was preceded by the same valence or a contrasting valence. This results in two groups: 1) both subjects experienced an event in the same context (i.e. *r*_(A1,A1)_. *r*_(B1,B1)_, *r*_(A2,S2)_, or *r*_(B2,B2)_) 2) both subjects experienced an event in different contexts (i.e. *r*_(A1,A2)_ or *r*_(B1,B2))_. We then calculated the mean of the across-subject correlations and averaged across events within each grouping. We tested whether the average cross-subject pairwise correlations for emotional events that were heard in the same context were significantly greater than those heard in different contexts^28^:

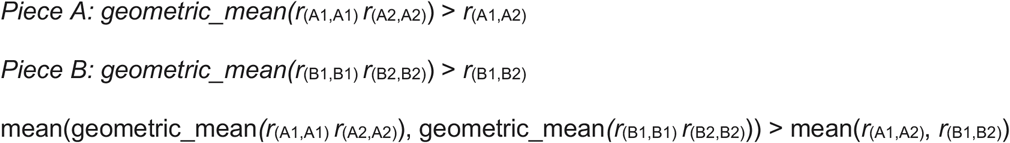

To assess statistical significance, we performed resampling analysis wherein the participant labels (whether each participant heard version 1 or version 2 of the pieces) are randomly shuffled 1000 times. For each shuffling, the cross-participant correlation matrix was recalculated and the means of cross-subject correlations binned according to the same groupings as above (now randomized) were re-calculated. The true difference in correlation values was compared to the null distribution in which context is random to produce a p-value for every region/searchlight. Resulting statistical maps were cluster-thresholded (cluster-size at p value < 0.05).

#### Systematic shifts in timing of transitions based on context

To evaluate how preceding context influences the timing of emotional experience in the brain, we tested whether systematic shifts in the timing of event transitions occurred when the event was preceded by an event with a similar or contrasting valence, using a previously-developed approach^30^. A measure of the “speed” of the transition from one emotion to the next can be obtained using the probability function of the HMM, which, for each time point, gives a value (between 0 of 1) of the likelihood that the brain is in a particular state. For each emotional event, we selected in both version A and version B the timepoints corresponding to this event and its preceding event. Because the preceding event differs across the two conditions, we cropped the longer one to have the same length as the shorter one, ensuring that the number of timepoints was matched across conditions. We ran separate HMMs for group average data for these timepoints for each searchlight/ROI, setting the number of events in the model to 2. We then calculated the expected value of the event assignment at each timepoint (dot product of the probability function with event labels) and summed this expected value across timepoints (higher sums correspond to faster transitions; see Fig. 6A). The difference in this sum between the two conditions indicates the number of timepoints by which the event transition is shifted, which can be converted into units of seconds by multiplying by the TR (1.5s).

To compare how the preceding emotional event changes the speed of transition we categorized all events across the two pieces into one of 4 conditions: a positive emotional event preceded by a negative emotional event (NP), positive emotion preceded by a positive emotion (PP), negative emotion preceded by a positive emotion (PN), and negative emotion preceded by a negative emotion (NN). A t-statistic (mean divided by standard deviation) was calculated to compare differences in speed of transition between the NN and PN condition and then PP and NP conditions and the two resulting values were averaged. Statistical significance of the averaged t-statistic was evaluated through bootstrapping, i.e. by randomly sampling data from the 40 participants with replacement 1000 times, re-running the above analysis and re-calculating the test statistics.

## Data availability

Simulated data used for power analysis are made accessible for peer-review of the Stage 1 manuscript. Upon acceptance of the manuscript, fMRI images in BIDS format will be published on OpenNeuro. We will also publish the musical stimuli and behavioral ratings as a dataset to be used by researchers interested in music and emotions.

## Code availability

Code used to conduct the all analyses are made available on OSF (https://osf.io/a57wu/?view_only=996150c5d6fc49c1a004c78ef2f852d3). Upon acceptance of the manuscript, all code will be made publicly available on OSF as well as the first author’s Github page, so that independent researchers reproduce the results.

## Acknowledgements

This project was funded by internal research funding by the Center for Science and Society at Columbia University, awarded to the first author. The funders have/had no role in study design, data collection and analysis, decision to publish or preparation of the manuscript.

## Author contributions

M.S., K.O. and C.B. conceived and designed the study. M.S. developed the online rating system. M.S. collected all data, wrote most of the code, and wrote the manuscript. C.B. provided methodological advice and provided some of the code. K.O. provided theoretical advice and assisted with manuscript preparation.

