## Supplementary Information for "Brain state dynamics reflect emotion transitions induced by music"

### Supplementary Materials

#### Additional retrospective recall analyses

In addition to timing, we tested if the intensity of post-listening emotions ratings varied by emotional context, i.e. if the version of the piece that a person heard systematically altered how people remembered those moments within the musical piece. If self-report ratings varied by piece version, this would suggest that the proceeding emotional context is influencing how the music is being experienced and providing strong motivation for Aim 3 of our proposed fMRI analysis. Repeated measures ANOVAs (*rm_anova* function from the Pingouin Python package ^74^) were used to determine if ratings (vividness, surprise, emotional intensity) varied by version of the piece and emotional label. All post-hoc comparisons were made with Tukey’s Honestly significant difference test (HSD) and significant results after multiple-comparison correction are presented below.

For *vividness* ratings, joy clips that were preceded by calm clips were remembered more vividly than when preceded by anxious clips (M_calm_ = 5.00, M_anxious_ = 4.46, p-value = 0.004) and sad clips (M_sad_ = 4.48, p-value = 0.043). In addition, sad clips preceded by calm clips were remembered more vividly than those preceded by joy (M_calm_ = 4.46, M_joy_ = 4.46, p-value = 0.016) or anxious (M_anxious_ = 3.87, p-value = 0.025). For *intensity* of felt-emotions ratings, joy clips that were preceded by anxious clips were rated as less joyful than those that were preceded by sad clips (M_anxious_ = 4.85, M_sad_ = 5.74, p-value = 0.024). For *surprise* ratings, sad clips that were preceded by calm were rated as more surprising than those preceded by anxious (M_calm_ = 3.06, M_anxious_ = 2.58, p-value = 0.04). Finally, for *enjoyment* ratings, anxious clips that were preceded by joyful clips were significantly less enjoyed than when preceded by calm (M_joy_ = 2.91, M_calm_ = 3.86, p-value = 0.001) or sad clips (M_sad_ = 3.51, p-value = 0.003).

#### Spatial correlations accounting for hemodynamic lag

It is possible that the spatial patterns that reflect differences in musical context that we observed are due to lag in the fMRI signal bleeding over from the preceding event. If this were the case, then it could be that the results simply reflect that the group that heard the piece in the same context (piece A vs. piece B) show more similar activation patterns for events because the preceding event was the same as well and not that the spatial signal in the auditory cortex is reflecting any change in context. To address this, we re-ran the analyses above, including only brain data from the second half of each musical, emotional event. In this way, we can ensure that the mean signal across the second half of the event would not include any spillover signal from the event before.

The results are largely similar when using only the second half of each event as compared to the entire event. Significant differences as a result of context are still observed in the left and right temporal lobe, including the primary and secondary auditory cortex and anterior temporal lobe. The results in the right precentral gyrus and sulcus, however, were no longer statistically significant.

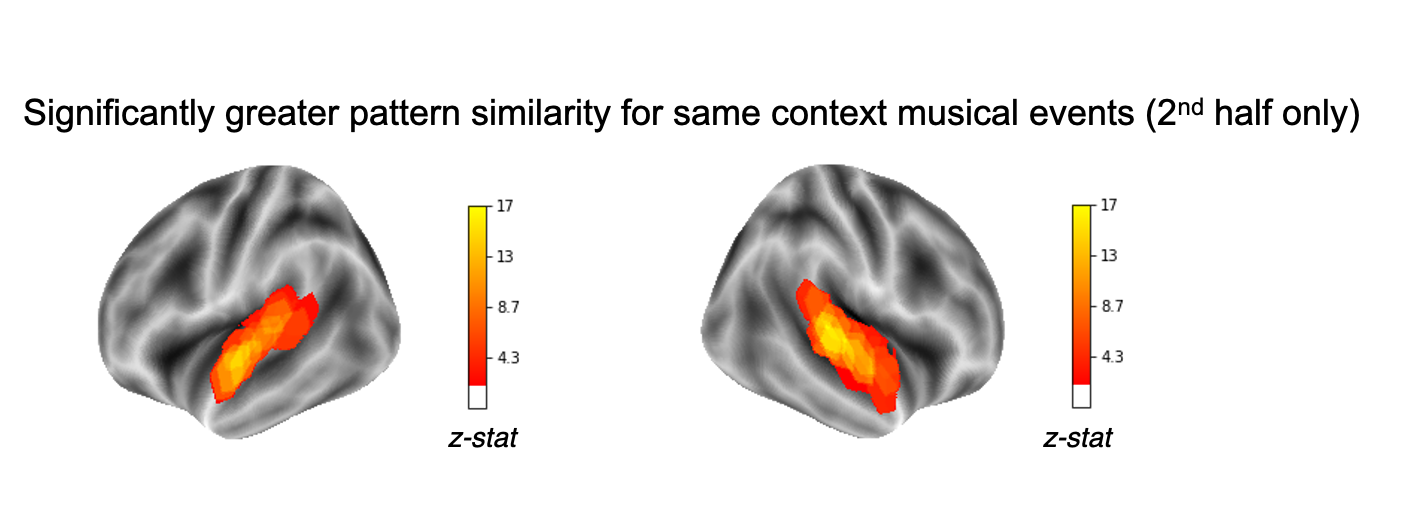

***Fig S1. Context-based changes in spatial brain patterns using only data from the second half of each emotional event****. Brain regions in which spatial patterns of the second half of each emotional event were significantly more similar in pairs of participants who heard them in the same condition (all in piece A or B, within group) as compared to pairs of participants who heard the music in different conditions (one in piece A other in B, across group). Colors correspond to z-stats of the ratio of across group vs. within group correlations as compared to a null model in which group membership was randomly permuted. Resulting statistical maps were cluster-corrected at p value < 0.05.*

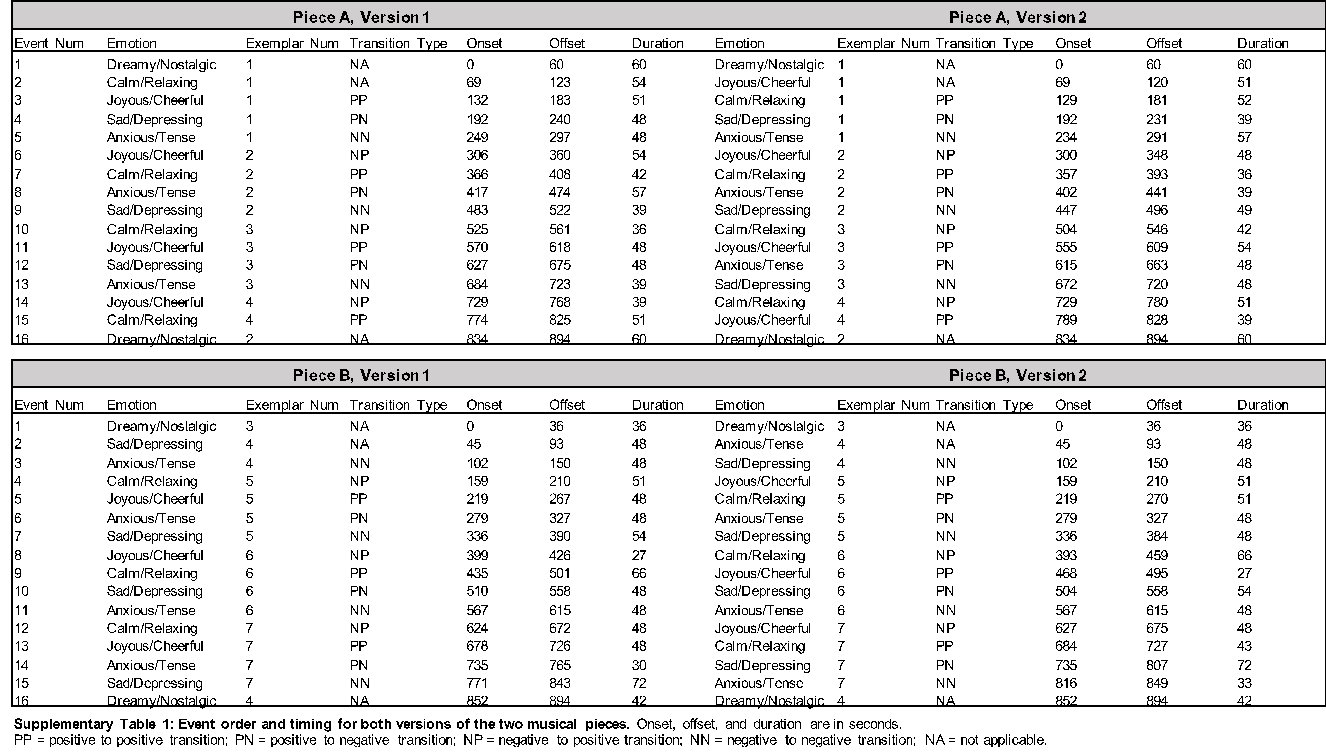
